# Superficial spinal Tac1-lineage neurons are polymodal nociceptive

**DOI:** 10.64898/2026.09.13.751298

**Authors:** Louison Brochoir, Pauline Larqué, Xinying Zhang, Charlotte Desnoyer, Artur Kania, Pascal Fossat, Yves De Koninck, Feng Wang

**Affiliations:** CERVO Brain Research Center, Quebec Mental Health Institute, Quebec City, QC, Canada; Graduate Program in Neuroscience, Faculty of Medicine, University Laval, Quebec City, QC, Canada; University Bordeaux, CNRS, IMN, UMR 5293, F-33000 Bordeaux, France; Montreal Clinical Research Institute (IRCM), Montreal, QC, Canada; Integrated Program in Neuroscience, McGill University, Montreal, QC, Canada; Sherbrooke University, Sherbrooke, QC, Canada; Division of Experimental Medicine, McGill University, Montreal, QC, Canada; Department of Anatomy and Cell Biology, McGill University, Montreal, QC, Canada; Department of Psychiatry and Neuroscience, University Laval, Quebec City, QC, Canada; The Center for Optics, Photonics and Lasers (COPL), University Laval, Quebec City, QC, Canada; Faculty of Dentistry, University Laval, Quebec City, QC, Canada

## Abstract

Tac1-lineage neurons in the spinal dorsal horn have been implicated in coping behavior in response to sustained noxious stimuli, but their developmental origin, cellular heterogeneity, and functional contribution remain incompletely understood. Here, we characterized the molecular identity, developmental trajectory, electrophysiological properties, and sensory responses of spinal Tac1-lineage neurons. In adults, Tac1-lineage neurons comprised a heterogeneous population of excitatory and inhibitory interneurons and excitatory projection neurons distributed across the superficial dorsal horn. In contrast, at embryonic day 16.5, Tac1-lineage neurons in laminae I/IIo were exclusively projection neurons, revealing a marked developmental transition in the composition of the Tac1 lineage. Electrophysiological recordings from spinal cord slices further demonstrated substantial functional heterogeneity, with most Tac1-lineage neurons exhibiting phasic or single-spike firing and smaller populations displaying tonic or reluctant firing. *In vivo* calcium imaging in anesthetized mice revealed prominent responses to noxious mechanical and thermal stimulation. Interestingly, Tac1-lineage neurons used distinct strategies to encode cold and heat intensity. Together, these findings reveal pronounced developmental and functional heterogeneity within the Tac1 lineage and demonstrate that adult Tac1-lineage neurons are predominately polymodal nociceptive.

## Introduction

The spinal dorsal horn is the first central site at which somatosensory information conveyed by primary afferents is integrated and transformed before being transmitted to the brain (Todd, 2010; Braz et al., 2014). It is organized into distinct laminae that differ in neuronal density, cellular composition, and patterns of primary afferent innervation (Todd, 2010). Most dorsal horn neurons are local interneurons, whereas projection neurons are concentrated in lamina I and distributed throughout the deeper laminae III-V (Todd, 2010; Wercberger and Basbaum, 2019; Graham and Hughes, 2020). Despite this well-defined anatomical organization, the extensive cellular and functional heterogeneity of dorsal horn neurons has complicated efforts to determine how specific spinal populations encode sensory information and generate appropriate behavioral responses.

Noxious stimuli evoke multiple behavioral outputs, ranging from rapid withdrawal reflexes to sustained coping responses, such as licking, guarding, and attending to an injured body region. These responses are thought to engage dissociable neural circuits and may reflect different components of nociceptive processing (Ma, 2022). Although ascending spinal pathways have been proposed to differentially contribute to sensory-discriminative and affective-motivational function (Wang et al., 2022), the spinal populations supporting these different outputs remain incompletely defined. Notably, genetic ablation of spinal Tac1-lineage neurons in mice markedly reduced licking and guarding elicited by persistent noxious stimulation while sparing rapid withdrawal reflexes (Huang et al., 2019), suggesting that Tac1-lineage spinal neurons are particular important for organized coping responses to ongoing noxious stimuli, which is a direct manifesto of the affective-motivational dimension of nociception. However, the sensory response properties and circuit mechanisms of these neurons remain poorly understood.

On the other hand, previous studies have primarily examined neurons expressing Tac1 in the adult spinal cord (Xu et al., 2013). Adult Tac1-expressing neurons represent approximately 20% of dorsal horn neurons (Davis et al., 2023) and comprise excitatory interneurons, projection neurons, and a smaller population of inhibitory interneuron (Ribeiro-da-Silva and Hokfelt, 2000; Gutierrez-Mecinas et al., 2017; Gutierrez-Mecinas et al., 2018). Tac1-expressing projection neurons are concentrated in layer I and scattered across deeper laminae (Cameron et al., 2015; Gutierrez-Mecinas et al., 2018; Polgar et al., 2020). They constitute approximately 40% of lamina I spinoparabrachial neurons (Polgar et al., 2020), and a subset projects to the medial thalamus (Polgar et al., 2020). Tac1-expressing excitatory interneurons include cells with extensive dendritic arborization and a delayed firing in response to electrical stimulation (Dickie et al., 2019), properties characteristic of radial neurons (Grudt and Perl, 2002). They receive monosynaptic excitatory input from local interneurons as well as Aδ and C primary afferent fibers, suggesting their recruitment during nociceptive stimulation (Grudt and Perl, 2002; Yasaka et al., 2010; Polgar et al., 2020). Consistently, Tac1-expressing neurons exhibit activity-marker expression (cFos and pERK) following noxious mechanical and thermal stimulation, as well as pruritic stimuli (Gutierrez-Mecinas et al., 2017), while suppression of their synaptic output impaired thermal nociceptive behavior (Polgar et al., 2020), further supporting a potential role for this population in pain processing.

Adult Tac1-expressing neurons, however, represent only a subset of the broader Tac1-lineage population. Importantly, the two populations also differ in their anatomical distribution. Tac1-lineage neurons are concentrated in the superficial laminae but are also distributed throughout the deeper dorsal horn, whereas adult Tac1-expressing neurons are more restricted to laminae□-□o (Gutierrez-Mecinas et al., 2017). Thus, the extent to which the physiological properties established for adult Tac1-expressing neurons can be generalized to the broad Tac1-lineage neurons remain uncertain. Moreover, the responses of Tac1-lineage neurons to natural sensory stimulation have not been systematically characterized under intact physiological conditions. Here, we combined *in vivo* calcium imaging during mechanical and thermal stimulation with *ex vivo* slice electrophysiology to characterize Tac1-lineage neurons in the superficial spinal dorsal horn. Our results reveal a functionally heterogeneous population preferentially recruited by noxious stimuli and demonstrate extensive integration of nociceptive information across sensory modalities.

## Methods

### Animals

All experiments were conducted in accordance with guidelines of the Canadian Council on Animal Care and were approved by the committee for animal protection of Laval university (CPAUL; protocols #2023-1038 and #2023-1171). Most experiments were performed on 6-8-week-old male and female mice. Animals were group-housed (2-5 mice per cage) in ventilated cages with *ad libitum* access to food and water under a 12-h light/dark cycle (lights on from 7:00 a.m. to 7:00 p.m.). Behavioral tests were performed between 9:00 a.m. and 2:00 p.m. in a climate-controlled room maintained at 23°C and 30-50% relative humidity.

*Tac1-Cre* mice (*B6;129S-Tac1^tm1.1(cre)Hze^/J*) were purchased from the Jackson Laboratory (#021877; JAX). They were used for viral injection and optogenetic manipulation of adult Tac1-expressing neurons or crossed with reporter lines for anatomical and functional studies of Tac1-lineage neurons. *Tac1-Cre* mice were crossed with *B6J.Cg-Gt(ROSA)^26Sortm96(CAG-GCaMP6s)Hze^/MwarJ* mice (#028866; Ai96, JAX) to generate *Tac1-GCaMP6s* mice for *in vivo* calcium imaging; *B6;Cg-Gt(ROSA)26Sor^tm14(CAG-tdTomato)Hze^/J* (#007914; Ai14, JAX), to generate *Tac1-tdTomato* mice for slice electrophysiology; or *B6;129-Gt(ROSA)26Sor^tm5(CAG-Sun1/sfGFP)Nat^/J* (#021039, JAX) to generate *Tac1-Sun1GFP* mice for anatomical studies. In these crosses, transient Tac1-driven Cre expression induces irreversible recombination of the reporter allele, resulting in permanent reporter expression in Tac1-lineage neurons even if Tac1 expression subsequently ceases.

### Spinal virus injection and optic fiber implantation

*Tac1-Cre* mice were deeply anesthetized with isoflurane in oxygen (4% for induction and 1.5-2% for maintenance). The back of the animal was shaved, and a midline skin incision was made to expose the musculature overlying the lumbar vertebrae. The muscles were gently separated to expose the T13/L1 vertebrae. The dorsal spinous process was removed, and a 1-mm-diameter hole was drilled through the left dorsal aspect of the vertebra. AAV2/1-CAG-Flex-ArchT-EGFP (#142-aav1, Canadian Neurophotonics Platform Viral Vector Core Facility) was injected at two rostrocaudal positions and two dorsoventral depths, for a total of four injections. At each site, 125 nL of virus was delivered at a constant rate of 100 nL/min. An optic cannula with a ceramic ferrule (#R-FOC-BL200C-39NA, RWD), cleaved to <0.5 mm, was inserted through the drilled opening and positioned in the space between the vertebral bone and spinal cord. The cannula was then secured to the vertebra with dental cement.

### Retrograde labeling from parabrachial nucleus

Six *Tac1-Sun1/sfGFP* mice were deeply anesthetized with isoflurane (4% for induction and 1.5% for maintenance). Cholera Toxin Subunit B (CTb) conjugated to Alexa Fluor 647 (#C34778; ThermoFisher) was reconstituted in phosphate buffered saline (PBS) at 1% (w/v) and injected bilaterally into the parabrachial nuclei (AP: -5.0 mm, ML: ±1.5 mm, DV: −2.75 mm relative to bregma). A maximum volume of 250 nL was delivered at each injection site at a constant rate of 100 nL/min. Five days after CTb injection, mice were transcardially perfused via the left ventricle with 4% paraformaldehyde in PBS.

### Behavior experiments

To assess coping behavior in response to sustained noxious thermal stimuli, mice were placed on a hot/cold plate (#BIO-CHP, Bioseb) and allowed to acclimate for 5 min at 32°C. Mice were then exposed to different test temperatures, with cutoff durations set as previously described by Huang et al.: 4 min at 46 °C, 3 min at 47 °C, 1 min at 50 °C, 30 s at 56 °C, and 5 min at 0 °C (Huang et al., 2019). Each temperature was tested without and with continuous light-induced inhibition (520 nm, 10 mA). A minimum interval of 15 min was allowed between temperature tests. Behavioral responses were video-recorded and manually scored using BORIS software, including paw licking, lifting and shaking, as well as jumping. An integrated nociception score was calculated by assigning 1 point for shaking, 2 points for lifting, 3 points for licking, and 4 points for jumping.

### In vivo Ca^2+^ imaging

*In vivo* Ca^2+^ imaging was performed as previously described (Wang et al., 2018). Briefly, *Tac1-GCaMP6s* mice were deeply anesthetized with ketamine (100 mg/kg), xylazine (15 mg/kg), and acepromazine (2.5 mg/kg) diluted in saline (Vrontou et al., 2013). A laminectomy was performed to expose the L4-L5 spinal segments, and the spinal cord was stabilized using a custom-made fixation device. A pool was formed around the exposed spinal cord using 3% agarose and filled with Ringer’s solution containing (in mM): 126 NaCl, 2.5 KCl, 2 CaCl2, 2 MgCl2, 10 D-Glucose, and 10 HEPES (pH = 7.0). To visualize the vasculature, mice received a retro-orbital intravenous injection of 1% Texas Red-conjugated dextran (70 kDa, neutral, D1830; Invitrogen) diluted in saline.

The animals, together with the spinal stabilization device, were mounted on a custom-build video-rate two-photon microscope. A tunable femtosecond laser (InSight X3, Spectra-Physics) was set to an excitation wavelength of 940 nm. Emission filters of 500-550 nm and > 605 nm were used to detect GCaMP6s and Texas Red signals, respectively. Images were acquired through a 40x water-immersion objective (Olympus), at a spatial resolution of 0.375 µm/pixel and a frame rate of 32 frames/second.

Thermal stimuli were delivered to the plantar side of the hind paw using a feedback-controlled Peltier device with a 1 cm by 1 cm thermal probe (TSA--NeuroSensory Analyzer, Medoc). Temperatures ranging from 6°C to 50°C were delivered using a rapid ramp-and-hold protocol. The rate of temperature increase and decrease was set to 8°C/s and 4°C/s, respectively. For heating stimuli, the baseline temperature was 25°C, and target temperature of 38°C, 43°C, and 50°C were maintained for 15, 10, and 5 s respectively. For cooling stimuli, the baseline temperature was 32°C, and target temperature of 20°C, 15°C, and 6°C were maintained for 15, 10, and 8 s respectively. Relative shorter durations of noxious heat and cold stimuli were selected to minimize sensitization and prevent tissue injury.

Innocuous brushing and mild noxious pinching of the hind paw were performed using a small paintbrush and serrated forceps, respectively. Each mechanical stimulation epoch consisted of repeated application lasting a total of 15 s for brushing) and 8 s for pinching. For both thermal and mechanical stimuli, an interval of 3 to 5 minutes was maintained between successive stimuli to minimize neuronal sensitization.

### Ca^2+^ imaging data analysis

Raw image sequences were converted to TIFF format using ImageJ. The resulting TIFF movies were registered using a custom MATLAB (MathWorks) function that applied rigid-body translation alignment based on two-dimensional cross-correlation to correct for motion. A rectangular region of interest (ROI) was drawn in an area containing no visible neuron and used to estimate background fluorescence. For each frame, the mean fluorescence intensity within this background ROI was subtracted from every pixel in the corresponding frame.

Neuronal ROIs were manually drawn within the cytoplasm of visually identified neurons. The fluorescence intensity at each time point (Ft) was calculated as the mean pixel intensity within the corresponding ROI. Ca^2+^ signals were expressed as Δ*F/F_0_* = (*Ft* - *F_0_*)/*F_0_*, where *F_0_* was the mean fluorescence intensity during the first 2 s of the recording. To prevent artifactual amplification caused by very small *F_0_* values in neurons with low basal fluorescence, the denominator was set to 1 when the calculated *F_0_* < 1; the original *F_0_* value was retained in the numerator. The extracted Ca^2+^ traces were then synchronized with the corresponding thermal or mechanical stimulation data. Image registration, background subtraction, signal extraction, and synchronization were performed using custom MATLAB functions.

Positive responses were detected and quantified automatically using a separate custom analysis tool written in Spike2 (Cambridge Electronic Design), which provided an integrated interface for to trace visualization and analysis. Ca^2+^ traces were first smoothed using a 1-s temporal window. The baseline period was defined as the interval beginning 1 s after the start of the recording and ending 1 s before stimulus onset, typically providing a baseline duration of 3-5 s. The mean (F_b_) and maximum (F_b-max_) Δ*F/F_0_* during this period were then calculated. A response was classified as positive if its peak Δ*F/F_0_* value during stimulation exceeded the following threshold:

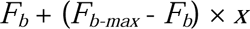

where *x* ranged from 2 to 3 depending on baseline stability. Responses lasting < 0.5 s were excluded because their duration was inconsistent with the relatively slow decay kinetics of GCaMP6s. Although the automated detection algorithm was highly reliable, all traces were also inspected visually to exclude false-positives and identify any false negatives.

Peak response amplitude was measured for each positive response obtained during repeated trials and then averaged. Data from all experiments were subsequently combined into a single database, and Microsoft Excel Pivot Tables were used to extract the appropriate subsets for statistical and cluster analyses.

### Spinal cord slice electrophysiological recordings

*Tac1-TdTomato* mice were deeply anesthetized with 30% urethane (0.1 mL/10g), and the vertebral column was exposed after removing the back skin. The lumbar segment of spinal cord was dissected and placed in a Sylgard silicone elastomer-coated Petri dish containing ice-cold cutting solution (in mM: 93 NMDG, 2.5 KCl, 1.2 NaH2PO4, 25 D-glucose, 20 HEPES, 30 NaHCO3, 5 sodium gluconate, 2 thiourea, 3 sodium pyruvate, 10 MgSO4, 0.5 CaCl2; pH 7.5-7.55; osmolarity 300-310mOsm), continuously oxygenated with carbogen (95% O2, 5% CO2). Residual nerve roots were cut and the meninges were removed, and the spinal cord was mounted on a vibratome stage in cold cutting solution. Parasagittal slices (350 µm) were prepared and incubated at 37°C for 15 minutes in the same solution.

After recovery, slices were transferred to and maintained in oxygenated recording solution (in mM: 92 NaCl, 30 NaHCO3, 25 D-glucose, 20 HEPES, 2.5 KCl, 1.2 NaH2PO4, 2 MgSO4, 2 CaCl2, 5 ascorbic acid, 2 thiourea, 3 sodium pyruvate, with 12 mM N-acetyl-L-cysteine; pH 7.3-7.4) at 37°C for an hour, before being transferred to room temperature.

Tac1-expressing neurons were identified using an Axioskop 2FS microscope (Zeiss) with a 530 nm excitation light to visualize tdTomato. Whole-cell patch-clamp recordings were performed using a borosilicate glass pipette (3-4MΩ) filled with the following intracellular solution (in mM): 5 K-methylsulfonate, 5 KCl, 2 MgCl, 10 HEPES, 4 NaATP, 0.4 NaGTP; pH 7.22; 290 mOsm. Neuronal firing properties and synaptic currents were assessed in current-clamp and voltage-clamp mode, respectively, using Multiclamp 700B amplifier (Molecular Devices) with a Digidata 1440 interface. Data were acquired at 20kHz and filtered at 10kHz with pClamp 10.3.

Cells were classified as tonic, phasic, single-spike, or reluctant based on their discharge patterns obtained in current-clamp mode at rest and in response to 12 steps of 500 ms current injections with 20 pA increment. Analysis of current-clamp and voltage-clamp data was performed using EasyElectrophysiology.

### Immunohistochemistry

Embryos were obtained from *Tac1-Sun1sf/GFP* mice following the protocol described by Roome (Roome et al., 2020). Pregnant females (16 days post-plug) were anesthetized via intraperitoneal injection of ketamine (100 mg/kg) and xylazine (15mg/kg) mixture. Embryos were dissected in cold 1xPBS, fixed in 4% paraformaldehyde (PFA) in PBS at 4°C for 2h with agitation, briefly washed and transferred to cryoprotective solution for 1-2 days before spinal cord dissection and sectioning. Adult mice were transcardially perfused with saline followed by 4% PFA containing 0.1% picric acid in phosphate buffer (pH 7.4). Spinal cords were dissected, post-fixed for 2h, and cryoprotected overnight in 30% sucrose. Embryonic spinal cords were cut at 20 µm using a cryostat (NX70, Thermo Fisher), whereas adult spinal cords were sectioned at 25 µm using a vibratome.

For immunostaining, sections were washed in PBS and permeabilized in PBS containing 0.3% of X-100 Triton (PBST) and either 1% bovine serum albumin (BSA) or 5% heat-inactivated horse serum (HIHS) for 30min. Sections were then incubated overnight at 4°C with primary antibodies (see Table 1) diluted in PBST containing 2% BSA or 1% HIHS. The following day, sections were washed and incubated for 1-2 hours at room temperature with secondary antibodies and a fluorophore-conjugated isolectin B4 for labeling. Sections were then mounted with either Dako mounting medium or 10% Mowiol/25% glycerol, and allowed to dry, and stored at 4°C.

### Confocal microscopy and image acquisition

Images were obtained with a confocal laser scanning microscope (LSM700, Leica) using either a 40x air objective (NA 1.4) or a 63x oil immersion objective (NA 1.4), at a resolution of 1024 x 1024 pixels and a pixel size of 0.066 µm.

### Statistical analysis

All statistical analyses were performed using GraphPad Prism (version 10.4.1). Two-tailed unpaired t-tests were used for comparisons between two groups, whereas comparisons among multiple groups were performed using one-way of two-way ANOVA, followed by post hoc tests as indicated in the figure legends. Data are presented as mean ± SEM, and differences were considered statistically significant at p < 0.05.

## Results

### Optogenetic inhibition of adult spinal Tac1-expressing neurons does not affect coping behavior

A previous study showed that ablation of spinal Tac1-lineage neurons almost completely abolished coping behavior in response to sustained noxious stimuli while leaving withdrawal reflexes intact (Huang et al., 2019). To determine whether acute inhibition of adult Tac1-expressing neurons produces a similar effect, we injected a Cre-dependent viral vector encoding the inhibitory opsin ArchT (Han et al., 2011) into the spinal cord of adult *Tac1-Cre* mice, thereby selectively expressing ArchT in adult Tac1-expressing spinal neurons. We then implanted an optical cannula on the left spinal cord to deliver green light (wavelength = 520 nm) for optogenetic inhibition.

Because Tac1 is also expressed in primary sensory neurons (Fig. S1), we first examined whether intraspinally injected virus could undergo retrograde transport to dorsal root ganglion (DRG) neurons. We injected a Cre-dependent GFP virus into the spinal cord of *Tac1-tdTomato* mice. GFP expression was observed selectively in spinal cord neurons (Fig. S2*a,b*), but was absent from DRG neurons (Fig. S2*c*), indicating that the injected virus was not detectably transported to primary sensory neurons and our optogenetic manipulation selectively targeted spinal Tac1-expressing neurons.

We next assess behavioral responses to sustained noxious cold and heat using cold- and hot-plate assays with and without optogenetic inhibition. Mice were video-recorded, and coping behaviors, including licking, shaking, lifting, and jumping, were analyzed offline. Surprisingly, optogenetic inhibition of adult spinal Tac1-expressing neurons did not alter either the number of licking episodes or total licking duration across the cold- and hot-plate temperatures tested (Fig. 1*a,b*). Similarly, shaking, lifting, and jumping behaviors were unaffected by optogenetic inhibition (Fig. 1*c*, S3). We further calculated a composite nociception score integrating these coping behavior and found no significant difference between trials with and without optogenetic inhibition (Fig. 1*d*).

**Figure 1.**
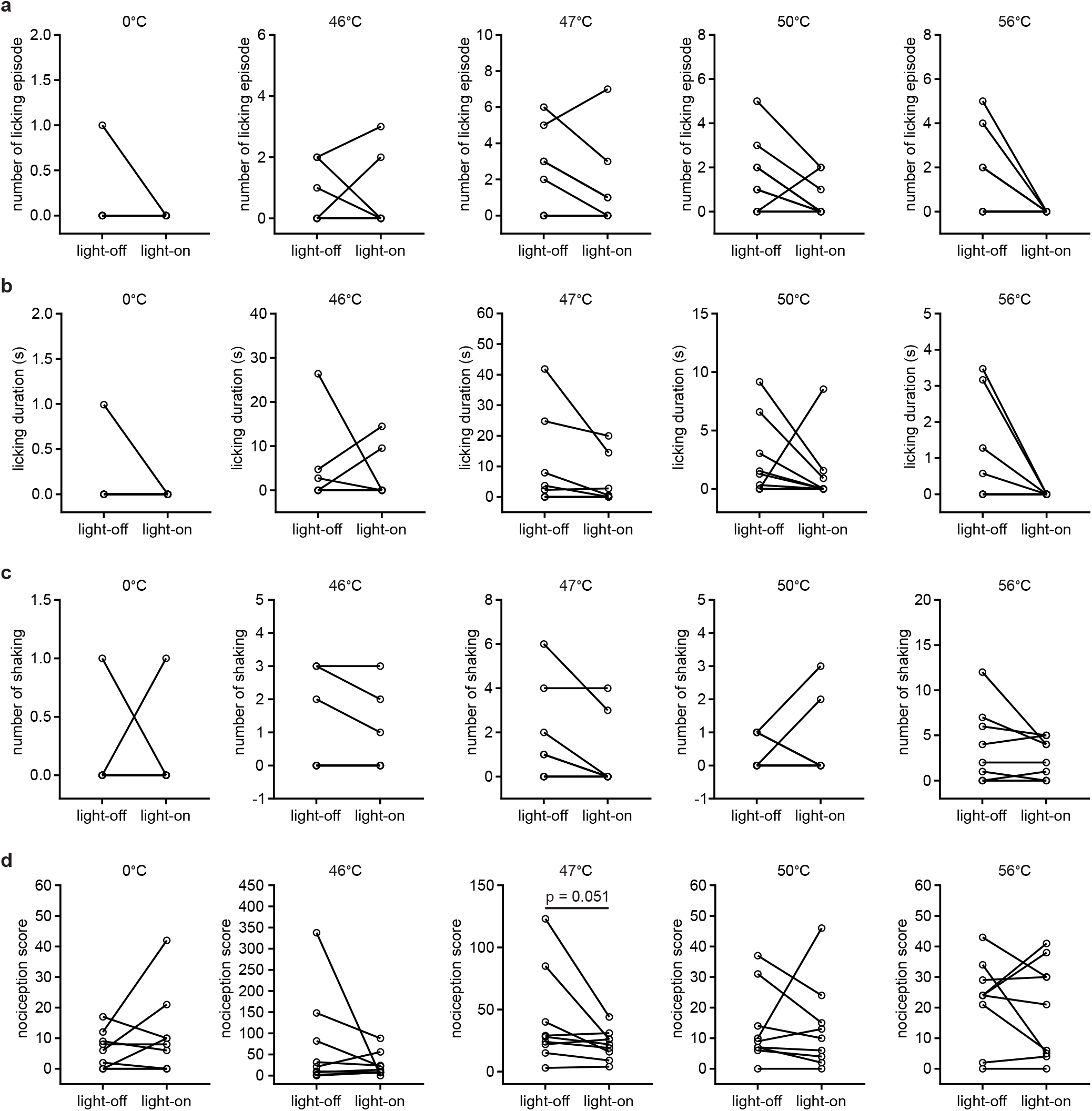
Adult spinal Tac1-expressing neurons are dispensable for coping behaviors evoked by sustained noxious thermal stimuli. Optogenetic inhibition of spinal Tac1-expressing neurons did not significantly alter the number of licking episodes (a), licking duration (b), number of shaking episodes (c), or integrated nociception score (d) evoked by hot- or cold-plate stimulation. The nociception score was calculated by assigning 1 point for shaking, 2 points for lifting, 3 points for licking, and 4 points for jumping. Statistical comparisons were performed using Wilcoxon matched-pairs signed-rank test. n = 9 *Tac1-Cre* mice.

Together, these results demonstrate that acute inhibition of adult spinal Tac1-expressing neurons does not significantly affect coping behavior evoked by sustained noxious thermal stimuli. In contrast to the pronounced behavioral effects previously observed following ablation of Tac1-lineage neurons, these findings suggest that Tac1-lineage neurons and adult Tac1-expressing neurons may represent functionally distinct populations.

### Spinal Tac1-lineage neurons comprise a heterogeneous population

Although spinal Tac1-lineage neurons have previously been described to include interneurons and projection neurons (Huang et al., 2019; Choi et al., 2020), accurately defining their cellular composition has been challenging. Previous studies largely relied on *Tac1-tdTomato* mice, in which tdTomato is also expressed in primary afferents (Fig. S1, S4). Dense labeling of Tac1-lineage afferent fibers in the dorsal horn could obscure tdTomato^+^ spinal neurons and complicate their identification (Fig. S4). To overcome this limitation, we generated *Tac1-Sun1GFP* mice, in which nuclear-localized GFP (Mo et al., 2015) selectively labels Tac1-lineage neurons (Fig. 2), thereby avoiding confounding signal from Tac1-lineage primary afferent fibers.

**Figure 2.**
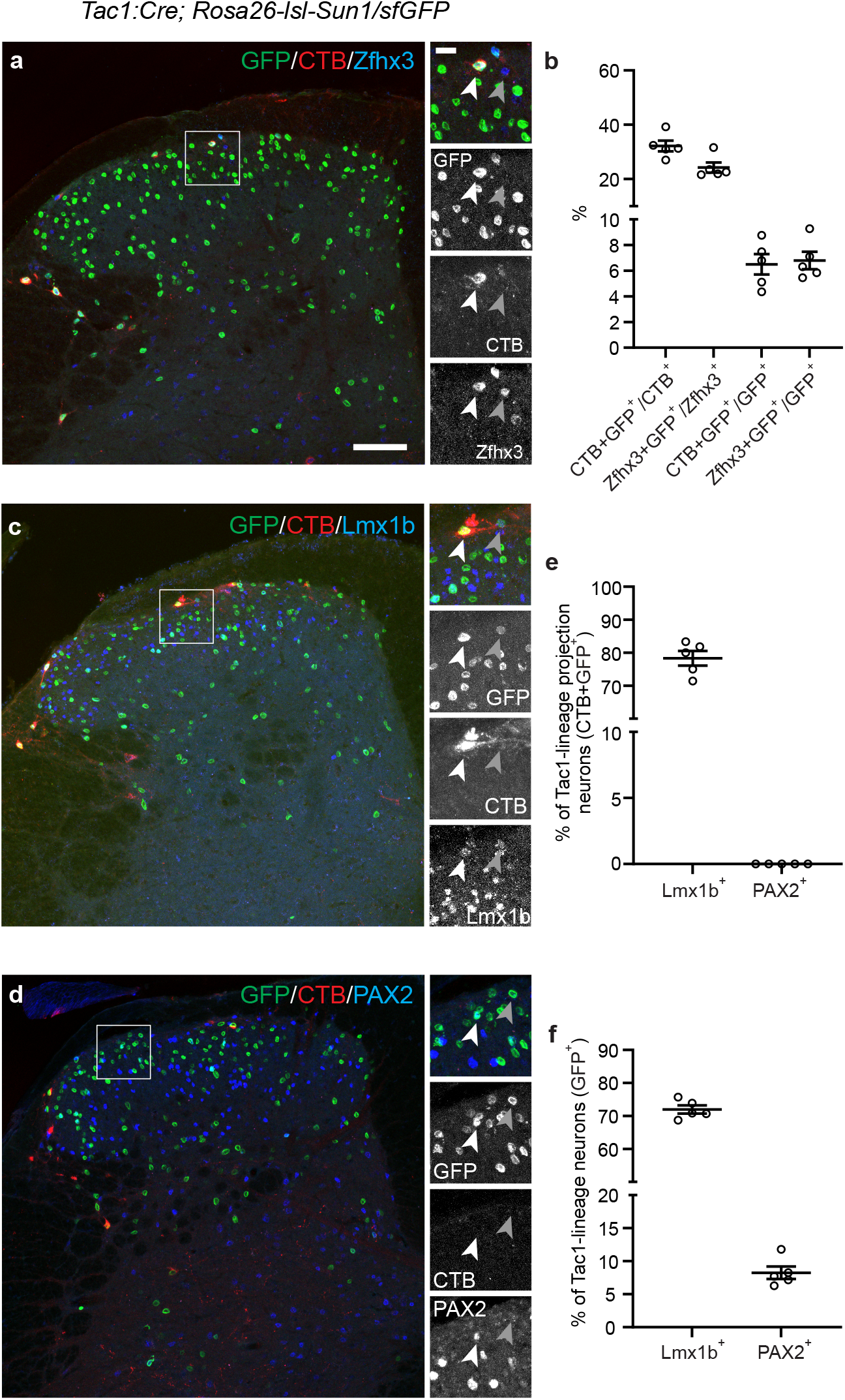
Spinal Tac1-lineage neurons comprise a heterogeneous population in adulthood. (a) Left: representative spinal cord section from a *Tac1-Sun1/sfGFP* mouse following CTb injection into the lateral parabrachial nucleus, showing GFP (green), CTb (red), and the projection neuron marker Zfhx3 (blue) in the dorsal horn. Right: higher-magnification images of the boxed region in the left panel, showing a GFP^+^/CTB^+^/Zfhx3^+^ Tac1-lineage projection neurons (white arrowhead) and a Zfhx3^+^/Tac1-negative projection neuron (gray arrowhead). Scale bars = 100 μm (left) and 20 μm (right). (b) Quantification of the overlap between Tac1-lineage and projection neurons in the superficial dorsal horn. Approximately 30% of projection neurons belonged to the Tac1-lineage, whereas only 7% of Tac1-lineage neurons were projection neurons (n = 5 *Tac1-Sun1/sfGFP* mice). (c) Left: representative spinal cord section from a *Tac1-Sun1/sfGFP* mouse following CTb injection into the lateral parabrachial nucleus, showing GFP (green), CTb (red), and the excitatory neuronal marker Lmx1b (blue). Right: higher-magnification images of the boxed region, showing a GFP^+^/CTB^+^/Lmx1b^+^ Tac1-lineage projection neuron (white arrowhead) and a GFP^+^/Lmx1b^+^ Tac1-lineage neuron without CTb labeling (gray arrowhead). (d) Left: representative spinal cord section from a *Tac1-Sun1/sfGFP* mouse following CTb injection into the lateral parabrachial nucleus, showing GFP (green), CTb (red), and the inhibitory neuronal marker Pax2 (blue). Right: higher-magnification images of the boxed region, showing a GFP^+^/Pax2^+^ Tac1-lineage inhibitory neuron (white arrowhead) and a GFP^+^/Pax2^-^ Tac1-lineage neuron (gray arrowhead). (e-f) Quantification of excitatory and inhibitory neuronal identities within the Tac1 lineage. (e) Most Tac1-lineage projection neurons were excitatory (Lmx1b^+^), whereas none were inhibitory (Pax2^+^). (f) Among the overall Tac1-lineage population, the majority (70%) were excitatory (Lmx1b^+^) and approximately 10% were inhibitory (Pax2^+^).

To determine the contribution of Tac1-lineage neurons to ascending spinal pathways, we injected the retrograde tracer cholera toxin subunit B (CTb) into the parabrachial nucleus of *Tac1-Sun1GFP* mice and combined retrograde tracing with immunohistochemistry against Zfhx3, a marker enriched in spinal projection neurons (Osseward et al., 2021). In laminae I and outer lamina II (IIo), 31% of CTb-labeled spinoparabrachial neurons were GFP^+^, indicating that they belonged to the Tac1 lineage (Fig. 2*a,b*). Among the overall population of Zfhx3^+^ projection neurons in laminae I/IIo, 22% were Tac1-lineage neurons (Fig. 2*a,b*). Conversely, only 5% of Tac1-lineage neurons in these laminae were projection neurons (Fig. 2*a,b*), confirming that the vast majority of Tac1-lineage neurons in laminae I/IIo are interneurons despite their substantial contribution to ascending spinoparabrachial output.

To determine the excitatory/inhibitory composition of this interneuron-dominated population, we examined expression of the excitatory neuron marker Lmx1b (LIM homeobox transcription factor 1-beta) (Ding et al., 2004; Borromeo et al., 2014) and the inhibitory neuron marker Pax2 (Paired Box 2) (Larsson, 2017; Das Gupta et al., 2021) in *Tac1-Sun1GFP* spinal cords. As an internal validation of these markers, we first examined Tac1-lineage spinoparabrachial projection neurons, whose excitatory identity is well established. As expected, 78% of Tac1-lineage spinoparabrachial neurons expressed Lmx1b, whereas none expressed Pax2 (Fig. 2*c-e*), consistent with the predominantly excitatory nature of ascending spinal projection neurons (Littlewood et al., 1995; Cameron et al., 2015; Wercberger and Basbaum, 2019). This confirmed that our Lmx1b/Pax2 labeling reliably distinguishes excitatory from inhibitory identity.

We next examined the entire Tac1-lineage population in laminae I/IIo. Among these neurons, 72% expressed Lmx1b, whereas only 7% expressed Pax2 (Fig. 2*c,d,f*), consistent with previous reports (Huang et al., 2019; Choi et al., 2020). Thus, although the adult Tac1-lineage is predominantly excitatory, it also contains a small inhibitory population.

Together, these results demonstrate that at adult stage, Tac1-lineage neurons in laminae I/IIo: comprise a heterogeneous population consisting predominantly of excitatory interneurons, together with a small population of inhibitory interneuron and excitatory projection neurons. This marked heterogeneity raises an important developmental question: are these distinct Tac1-lineage populations specified together early in development, or do different populations acquire Tac1 expression at distinct developmental stage? To address this question, we next examined the Tac1-lineage in the embryonic spinal cord.

### Embryonic Tac1-lineage neurons are exclusively projection neurons

To determine when Tac1 expression emerges and which neuronal populations initially belong to the Tac1 lineage, we examined *Tac1-Sun1GFP* spinal cords at embryonic day 16.5 (E16.5), and co-labeled sections for the projection neuron marker Zfhx3 (Osseward et al., 2021) and Ebf1 (Early B-Cell Factor 1), a marker of differentiated, post-migratory neurons (Garel et al., 1999). At this stage, Tac1-lineage neurons were concentrated in lamina I and the deep dorsal horn (Fig. 3), in striking contrast to their adult distribution, in which Tac1-lineage neurons are predominantly located in lamina II (Fig. 2) (Huang et al., 2019; Choi et al., 2020).

**Figure 3.**
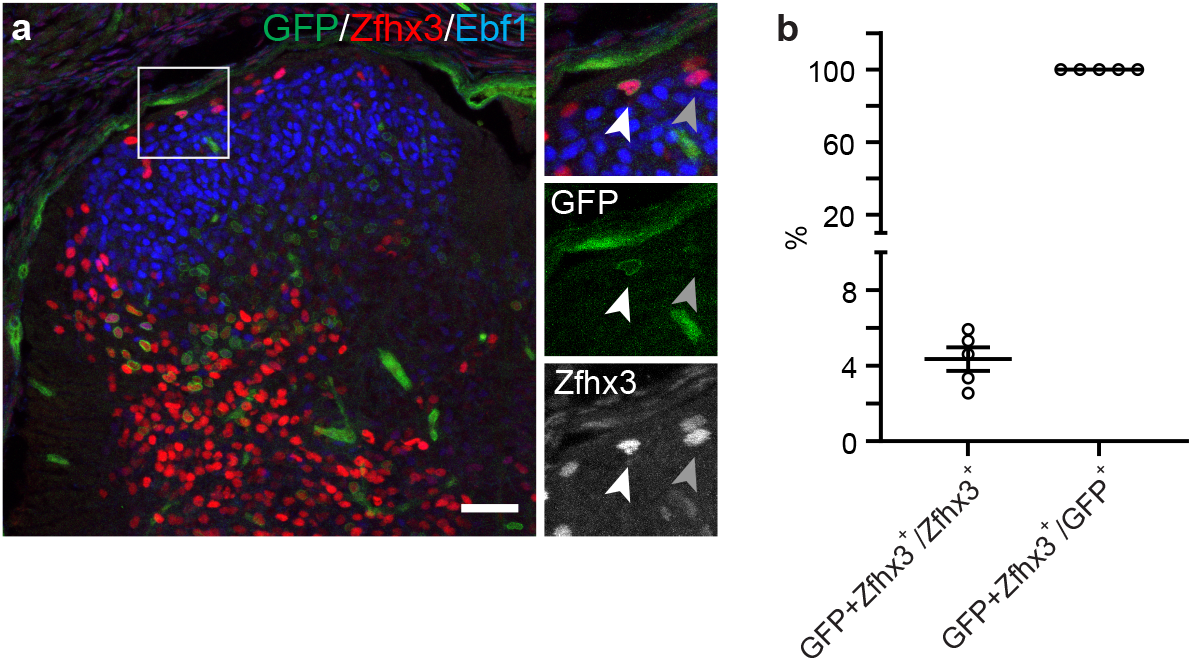
Tac1-lineage neurons in the superficial dorsal horn are exclusively projection neurons at E16.5. (a) Left: representative spinal cord section from an E16.5 *Tac1-Sun1/sfGFP* mouse, showing GFP (green), the projection neuron marker Zfhx3 (red), and post-migratory neuronal marker Ebf1 (blue). Right: higher-magnification images of the boxed region in the left panel, showing a GFP^+^/Zfhx3^+^ Tac1-lineage neuron (white arrowhead) and a Zfhx3^+^/GFP^-^ neuron (gray arrowhead). Scale bars = 100 μm (left). (b) Quantification of Tac1-lineage and Zfhx3^+^ neurons in the superficial dorsal horn. Virtually all Tac1-lineage neurons were Zfhx3^+^, consistent with a projection-neuron identity, whereas Tac1-lineage neurons accounted for < 5% of the total Zfhx3+ projection-neuron population (n = 5 *Tac1-Sun1/sfGFP* mice).

This difference in laminar distribution was accompanied by a striking difference in cellular composition. In laminae I/IIo, approximately 4% of projection neurons belonged to the Tac1-lineage, and all Tac1-lineage neurons identified in this region expressed the projection neuron marker Zfhx3 (Fig. 3). We found no evidence of Tac1-lineage interneurons in laminae I/IIo at E16.5 (Fig. 3). Importantly, Ebf1 expression confirmed that the majority of dorsal horn neurons in this region were already differentiated and post-migratory at this developmental stage (Fig. 3a). Thus, the absence of Tac1-lineage interneurons at E16.5 is unlikely to reflect a general lack of neuronal differentiation.

These findings reveal a striking developmental shift in the composition of the Tac1 lineage. Whereas Tac1-lineage neurons in adult laminae I/IIo are predominantly interneurons, all Tac1-lineage neurons detected in this region at E16.5 expressed Zfhx3 and exhibited a projection-neuron identity. Thus, the interneuron populations that predominate within the adult Tac1 lineage do not exhibit detectable Tac1 expression at E16.5.

### Tac1-lineage neurons predominantly exhibit phasic and single-spike firing

Having established the anatomical and developmental heterogeneity of the Tac1 lineage, we next asked whether Tac1-lineage neurons also exhibit diverse intrinsic electrophysiological properties. We prepared parasagittal spinal cord slices using *Tac1-tdTomato* mice and recorded from tdTomato^+^ neurons in laminae I–II (Fig. 4*a*). In current-clamp mode, the membrane potential was maintained at approximately −65 mV by injecting bias current, and neurons then were subjected to a series of twelve 500-ms current steps ranging from −20 pA to 200 pA in 20 pA increments. Their responses were used to classify their firing profile (Fig. 4*b*).

**Figure 4.**
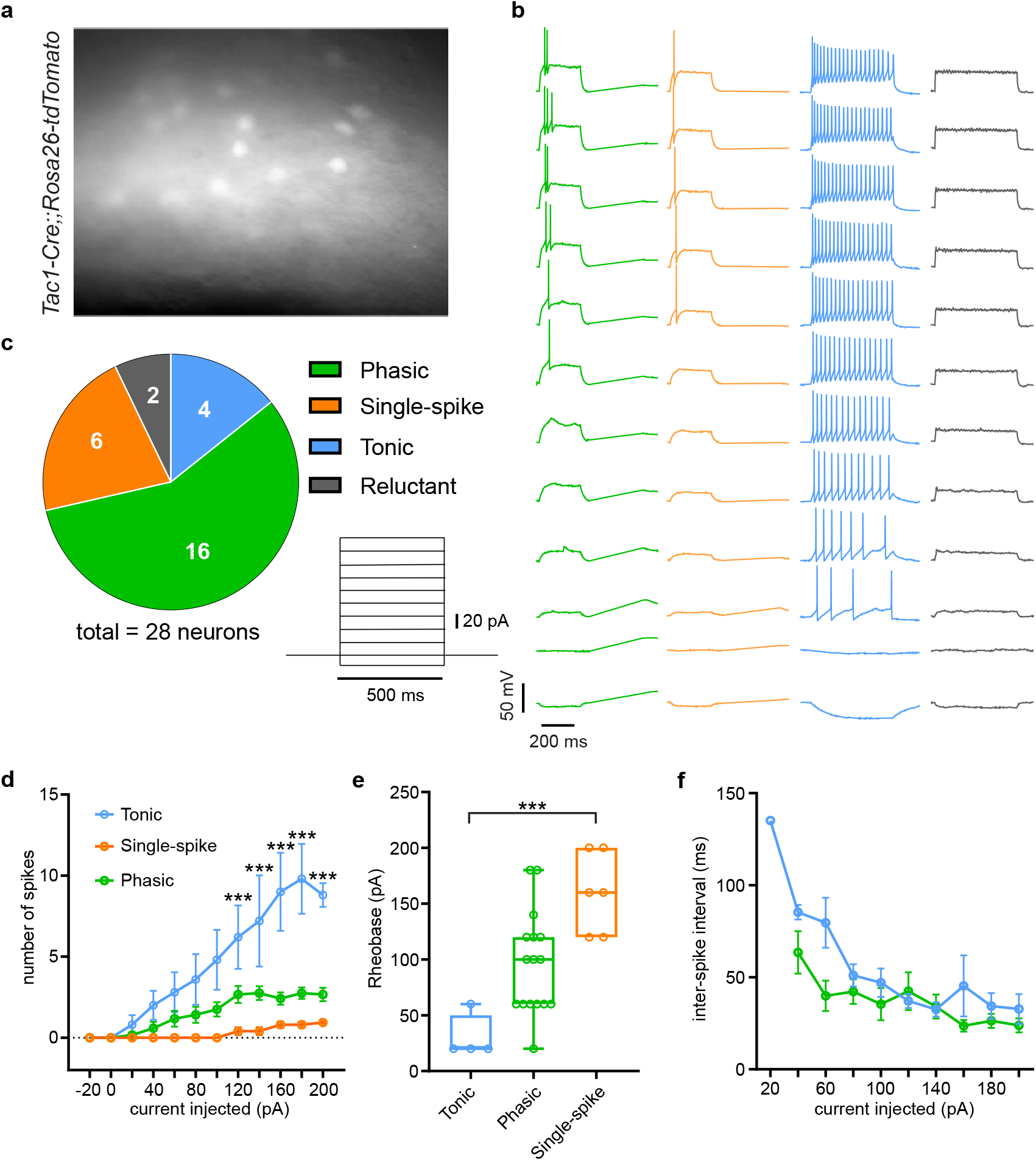
The majority of spinal Tac1-lineage neurons exhibit transient firing patterns. (a) Representative parasagittal spinal cord slice from a *Tac1-tdTomato* mouse showing Tac1-lineage neurons in the superficial dorsal horn. (b) Representative action potential firing patterns of Tac1-lineage neurons in response to current injections. Neurons were classified into four electrophysiological types: phasic, characterized by a high-frequency burst of variable duration; tonic, characterized by regular firing throughout the depolarizing current step; single-spike, characterized by a single action potential even in response to strong depolarization; and reluctant, characterized by a failure to generate action potentials within the range of current injections tested. For all recordings, the membrane potential was maintained at approximately -65 mV before current injection. (c) Distribution of firing patterns among superficial Tac1-lineage neurons. The majority exhibited transient firing patterns, comprising the phasic and single-spike types. (d) Tonic neurons generated significantly more action potentials than phasic and single-spike neurons (***p < 0.001, mixed-effects two-way ANOVA followed by Tukey’s multiple comparisons test; n = 12 phasic, 5 single-spike, and 5 tonic neurons). (e) Tonic neurons exhibited a significantly lower rheobase than single-spike neurons (***p<0.001, Kruskal-Wallis test followed by Dunn’s multiple comparisons test; n = 16 phasic, 6 single-spike, and 4 tonic neurons). (f) Inter-spike intervals did not differ significantly between phasic and tonic neurons (mixed-effects two-way ANOVA test; n = 12 phasic and 5 tonic neurons).

We identified four firing subtypes that occurred at markedly different frequencies (Fig. 4*c*). Phasic neurons, which generated a brief burst of action potentials at stimulus onset, were the most common subtype (55%). Single-spike neurons, which generated only one action potential even as current intensity increased, constituted 21% of recorded neurons. Tonic neurons, which sustained firing throughout the current step, accounted for 17%, whereas reluctant neurons, which did not fire within the tested current range, comprised the remaining 7% (Fig. 4*b,c*). Thus, although Tac1-lineage neurons were electrophysiological heterogeneous, firing patterns with a transient onset response, including phasic and single-spike firing, accounted for the majority (76%) of the recorded population.

Tonic neurons generated more action potentials during current injection than the other subtypes (Fig. 4*d*) and exhibited a significantly lower rheobase than single-spike neurons (Fig. 4*e*; 52.0 ± 23.32 pA vs. 160 ± 14.61 pA), indicating a greater capacity for sustained repetitive firing. Interestingly, although tonic neurons generated more action potentials than phasic neurons (Fig. 4*d*), their inter-spike intervals were comparable across the tested current intensities (Fig. 4*f*). The principal difference between these firing patterns therefore appeared to be the duration of the spike train rather than the instantaneous firing frequency within the train.

Together, these results demonstrate that Tac1-lineage neurons in the superficial dorsal horn comprise an electrophysiologically heterogeneous population. Most exhibit phasic or single-spike responses, whereas a small tonic subpopulation shows a greater capacity for sustained repetitive firing.

### Action potential waveform and afterhyperpolarization properties further distinguish Tac1-lineage subtypes

We next asked whether the firing-pattern subtypes identified above also differ in their action potential properties. Action potential threshold differed significantly between tonic and single-spike neurons (−23.7 ± 2.65 mV vs. −34.47 ± 3.07 mV), but not between tonic and phasic neurons (Fig.5*a*). In contrast, action potential amplitude, duration, and depolarization rise time were comparable across the three subtypes (Fig. 5*b*-*e*). Thus, most properties associated with action potential initiation and depolarization were similar across Tac1-lineage firing subtypes, although single-spike neurons exhibited a more hyperpolarized action potential threshold than tonic neurons.

**Figure 5.**
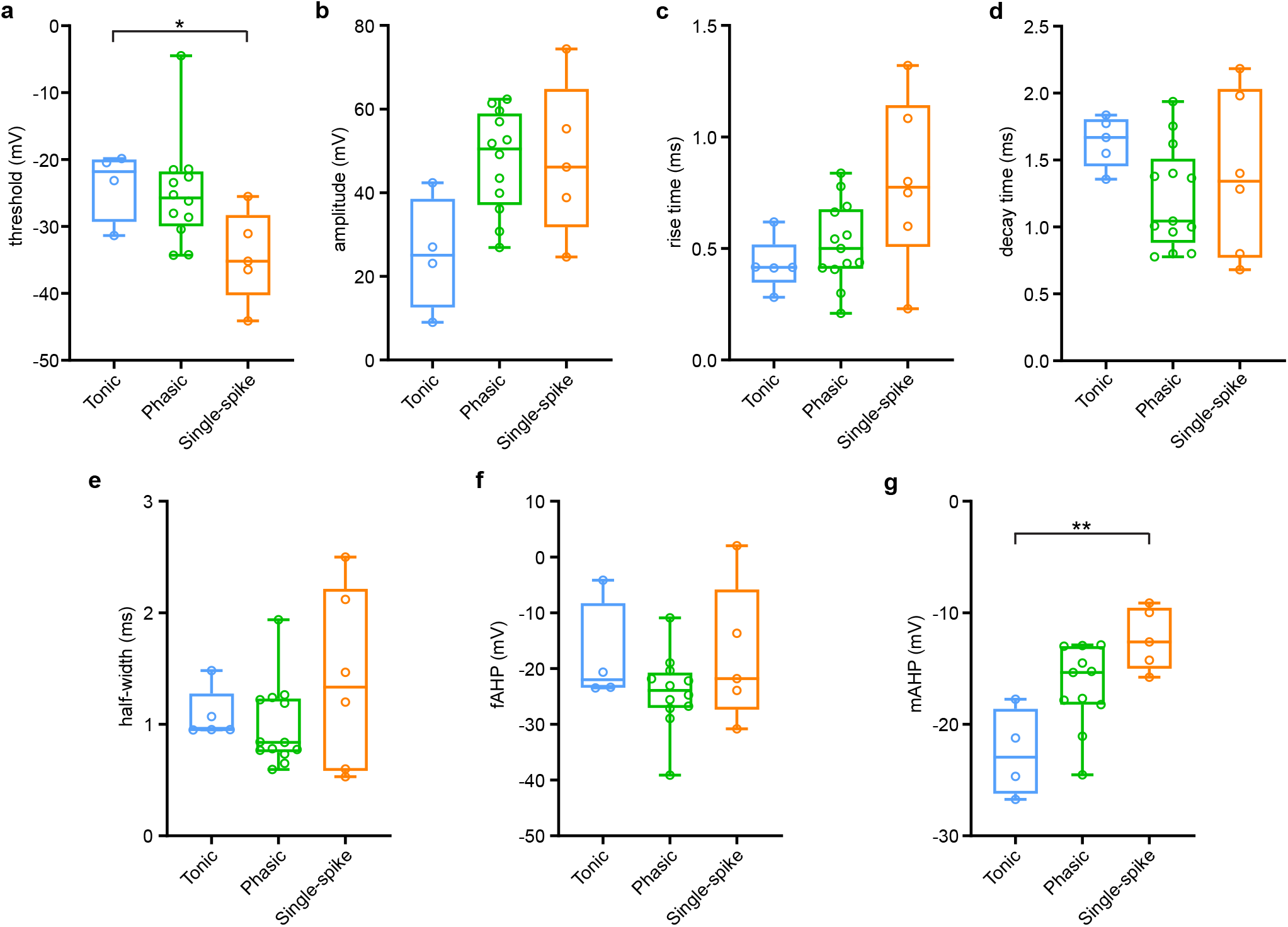
Active membrane properties of Tac1-lineage neurons. Quantification of action potential properties across different electrophysiological types of Tac1-lineage neurons, including action potential threshold (a), amplitude (b), rise time (c), decay time (d), half-width (e), fast afterhyperpolarization (fAHP; f), and medium afterhyperpolarization (mAHP; g). Tonic neurons exhibited a significantly higher action potential threshold and shorter mAHP than single-spike neurons (*p<0.05 and **p<0.01, respectively; Kruskal-Wallis test followed by Dunn’s multiple comparisons test; n = 11-13 phasic, 5-6 single-spike, and 4-5 tonic neurons, depending on the parameter analyzed).

Subtype-dependent differences were also evident in the afterhyperpolarization (AHP). The fast AHP component (fAHP), which contributes to rapid membrane repolarization, was comparable across all subtypes (Fig. 5*f*). In contrast, the medium AHP (mAHP) was significantly larger in tonic neurons than in single-spike neurons (Fig. 5*g*; −22.59 ± 1.98 mV vs. −12.35 ± 1.25 mV). Thus, tonic neurons were distinguished from the other firing subtypes by a larger mAHP.

Together, these findings demonstrated that Tac1-lineage firing subtypes differ not only in their responses to sustained current injection but also in specific features. In particular, tonic neurons were distinguished by a larger mAHP, whereas single-spike neurons exhibited a more hyperpolarized action potential threshold than tonic neurons.

### *In Vivo*, superficial Tac1-lineage neurons respond to both thermal and mechanical stimuli

Previous studies using activity markers, such as cFos and pERK demonstrated that adult Tac1-expressing neurons can be activated by both mechanical and thermal stimulation (Gutierrez-Mecinas et al., 2017), while loss-of-function studies established an important role for this population in nociceptive processing (Polgar et al., 2020). However, the response properties of individual Tac1-lineage neurons across different stimulus modalities remains unclear. To address this question, we performed *in vivo* calcium imaging in *Tac1-GCaMP6s* mice.

Imaging in 5-7-week-old mice during thermal and mechanical stimulation of the hind paw revealed diverse response profiles among simultaneously recorded neurons. During thermal stimulation, some cells responded selectively to cooling or heating stimuli (e.g., neurons #2–#5), whereas others responded across a broad range of thermal stimuli (e.g., neuron #6; Fig. 6). In addition, majority of Tac1-lineage neurons responded to noxious mechanical pinching, whereas relatively few responded to innocuous brushing. These observations revealed substantial functional heterogeneity within the Tac1-lineage population and motivated a systematic population-level analysis of their sensory response properties.

**Figure 6.**
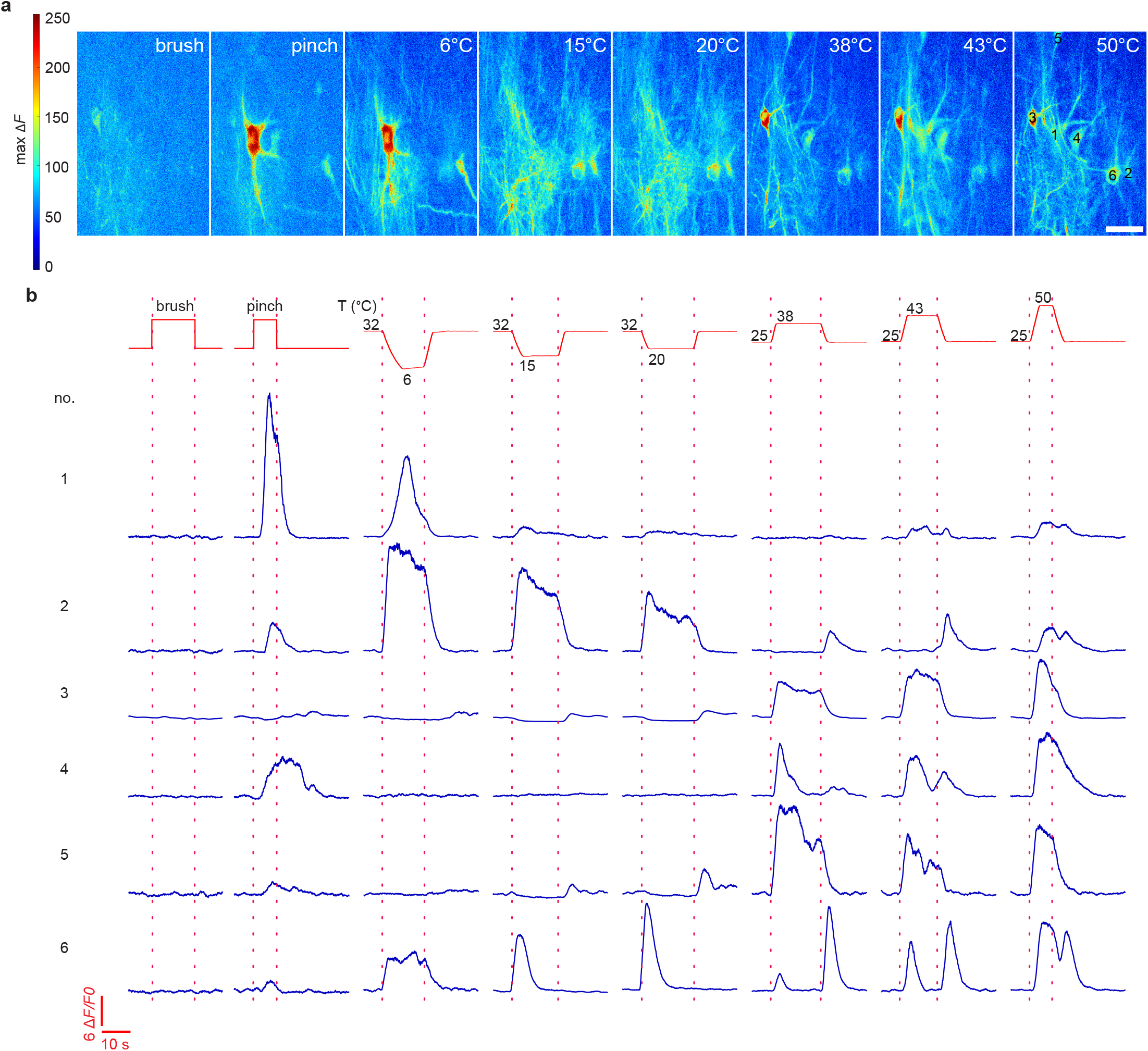
*In vivo* functional imaging of spinal Tac1-lineage neurons. (a) Color-coded heatmap of a representative imaging field showing the responses of individual Tac1-lineage neurons to innocuous and noxious thermal and mechanical stimuli. The color scale indicates the maximum ΔF response. (b) Representative Ca^2+^ curves from six neurons highlighted in (a), illustrating diverse response profiles to thermal and mechanical stimuli. Notably, neuron 6 responded both to heating and to the return phase following heating. Top: stimulus protocols for fast-ramp-and-hold thermal stimulation and the timing of manually applied brush and pinch stimuli. Baseline temperature of 32°C and 25°C were used for cooling and heating stimuli, respectively. Multiple repeated brush or pinch stimuli were applied during each mechanical stimulation trial.

### Superficial Tac1-lineage neurons are predominantly polymodal

We first quantified the proportion of Tac1-lineage neurons activated by each stimulus. Mechanically, innocuous brushing recruited relatively few neurons (<15%), whereas noxious pinching activated approximately 60% of the population. Thermally, 35–50% of neurons responded to cooling stimuli (6–20°C), whereas recruitment during heating increased with stimulus intensity: only ∼10% of neurons responded to innocuous warm (38°C), compared with more than 50% at the noxious heat (50°C; Fig. 7*a*). Together, these results indicate that Tac1-lineage neurons are preferentially recruited by noxious mechanical and thermal stimuli. No significant sex difference was observed (Fig. 7*a*), and data from males and females were therefore pooled for subsequent analyses.

**Figure 7.**
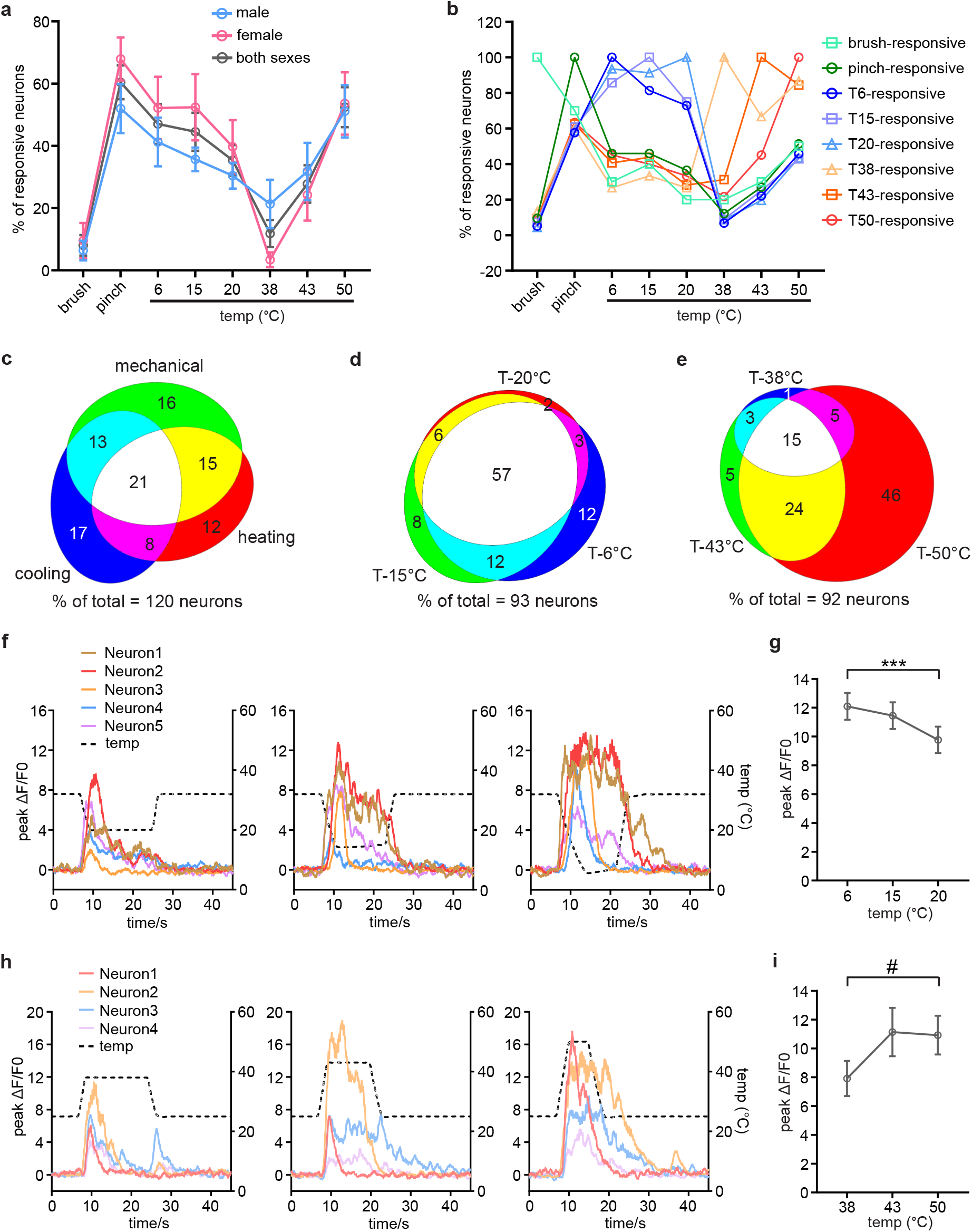
Spinal Tac1-lineage neurons use distinct strategies to encode cooling and heating. (a) Proportion of responsive neurons (defined as neurons responding to at least one type of stimulus applied to the hind paw) in each imaging field that responded to different mechanical and thermal stimuli. No significant sex differences were detected across stimuli (two-way repeated-measures ANOVA; n = 8 male and 9 female mice). Data are presented as mean ± SEM. (b) Proportion of neurons responsive to a given stimulus (indicated by different lines) that also responded to each of the other mechanical and thermal stimulus. (c) Euler diagram showing the overlap among mechano-responsive (responsive to brushing or pinching), heating-responsive (responsive to 38°C, 43°C, or 50°C), and cooling-responsive neurons (responsive to 20°C, 15°C, or 6°C). Numbers indicate the percentage of the total population of 120 mechano-responsive or thermo-responsive neurons. (d-e) Euler diagrams showing the overlap among neurons responsive to the three temperatures tested in the cooling (d) and heating (e) ranges. Numbers indicate the percentage of neurons responding to at least one of the three temperatures within the corresponding thermal range. (f) Representative Ca^2+^ traces from neurons responsive to all three cooling stimuli across the 20°C - 6°C range, corresponding to the white overlapping region in the Euler diagram in (d). (g) Peak response amplitudes of broad cool/cold neurons across cooling stimuli. Response amplitudes increased with cooling intensity and were significantly greater at 6°C than at 20C (***p < 0.001, Friedman test followed by Dunn’s multiple-comparisons test; n = 53 neurons). Data are presented as mean ± SEM. (h) Representative Ca^2+^ traces from neurons responsive to all three heating stimuli across the 38°C - 50°C range, corresponding to the white overlapping region in the Euler diagram in (e). (i) Peak response amplitudes of broad warm/heat neurons across heating temperature. Response amplitudes did not differ significantly with increasing temperature (Friedman test followed by Dunn’s multiple-comparisons test; n = 14 neurons). Data are presented as mean ± SEM.

We next examined the degree of overlap between stimulus modalities (Fig. 7*b*). Approximately 60% of neurons responsive to thermal or innocuous mechanical stimuli also responded to pinching. Similarly, 20–45% of neurons responsive to mechanical stimuli or heating temperatures (≥38°C) also responded to cooling (6–20°C). Among cooling-sensitive neurons, the likelihood of also responding to heating increased with stimulus intensity, from <10% at 38°C to >50% at 50°C. Within the cooling range, overlap was even greater, with 60–90% of neurons responding to multiple cooling temperatures (Fig. 7*b*). These data suggest extensive convergence of sensory modalities within individual Tac1-lineage neurons.

To quantify polymodality directly, neurons were classified as mechanosensitive, heating-sensitive, and/or cooling-sensitive according to their response profiles (Fig. 7*c*). Only 12–17% of neurons responded exclusively to a single modality. In contrast, more than 57% responded to at least two modalities, and approximately 21% responded to all three: a mechanical stimulus, a cooling stimulus (<20°C), and a heating stimulus (>38°C) (Fig. 7*c*). Thus, polymodal sensory integration, rather than modality selectivity, represents the predominant functional feature of superficial Tac1-lineage neurons.

### Tac1-lineage neurons use distinct strategies to encode cold and heat intensity

We next examined how Tac1-lineage neurons encode stimulus intensity within individual thermal modality. Among cooling-responsive neurons, approximately 80% responded to at least two of the three cold temperatures tested, and 57% responded to all three (Fig. 7*d*). Consistent with this broad responsiveness, lowering the temperature did not significantly increase the proportion of neurons recruited across the cooling range (Fig. 7*a*). In contrast, recruitment of heating-responsive neurons increased markedly with temperature (Fig. 7*a,e*). Among these neurons, 90% responded to the noxious 50°C stimulus, and 46% responded selectively to this temperature (Fig. 7*e*). Thus, increasing cooling intensity engages a largely overlapping population of Tac1-lineage neurons, whereas increasing heating intensity progressively recruits additional neurons.

We then asked whether response amplitudes of individual neurons varied with stimulus intensity. We compared calcium response amplitudes across temperatures within the cooling and heating ranges separately (Fig. 7*f*-*i*). Among cooling-responsive neurons, response amplitude increased significantly as temperature decreased, from 9.77 ± 0.91 (ΔF/F0) at 20°C to 12.09 ± 0.93 at 6°C (Fig. 7*f,g*). Thus, although stronger cooling did not recruit additional neurons, individual Tac1-lineage neurons increased their response amplitude with cooling intensity. In contrast, response amplitudes of heating-responsive neurons did not differ significantly across the temperatures tested (Fig. 7*h,i*), despite the progressive recruitment of additional neurons at higher temperatures.

Together, these results reveal two distinct strategies by which Tac1-lineage population encode thermal intensity. Within the cooling range, intensity is encoded primarily through amplitude scaling within a relatively stable neuronal population, whereas within the heating range, intensity is encoded primarily through progressive recruitment of additional neurons without substantial changes in the response amplitude of individual neurons.

## Discussion

Here, we show that superficial Tac1-lineage neurons are developmentally, electrophysiologically, and functionally heterogeneous. In adulthood, they comprise predominantly excitatory interneurons together with smaller inhibitory and projection-neuron populations, and most exhibit transient firing patterns. *In vivo*, Tac1-lineage neurons preferentially respond to noxious stimuli and encode cooling and heating intensity through distinct population strategies.

### Tac1-lineage neurons comprise a broader population than adult Tac1-expressing neurons

An important consideration is the distinction between Tac1-lineage neurons and neurons that actively expressing Tac1 in adulthood. In *Tac1-Cre* mice crossed with a Cre-dependent reporter line, Tac1-driven Cre expression induces irreversible recombination. Consequently, reporter-positive neurons include both cells currently expressing Tac1 and cells that expressed Tac1 earlier in development. Indeed, Huang et al. reported that only ∼45% of Tac1-lineage reporter-positive neurons continued to express detectable Tac1 mRNA (Huang et al., 2019), demonstrating that the lineage is considerably broader than the adult-expressing population. In contrast, approaches based on Tac1 mRNA, substance P expression, or viral targeting in adult *Tac1-Cre* mice more closely represent the adult Tac1-expressing population. This distinction is particularly relevant when comparing our results with previous studies from Todd and colleagues (Gutierrez-Mecinas et al., 2017; Dickie et al., 2019; Polgar et al., 2020).

Our adult anatomical data are broadly consistent with these studies. Approximately 72% of Tac1-lineage neurons in laminae / o expressed the excitatory marker Lmx1b, whereas only 7% expressed the inhibitory marker Pax2. Previous studies similarly identified adult Tac1-expressing neurons as predominantly excitatory, with a smaller (∼10%) inhibitory population (Gutierrez-Mecinas et al., 2017; Dickie et al., 2019). We further found that approximately 31% of spinoparabrachial neurons belonged to the Tac1 lineage. Nevertheless, projection neurons represented only a small fraction of the total Tac1-lineage population, indicating that in the adulthood, Tac1 lineage is dominated by interneurons.

Strikingly, this organization was different at E16.5, when virtually all Tac1-lineage neurons identified in laminae / o expressed the projection-neuron marker Zfhx3, whereas Tac1-lineage neurons accounted for only ∼4-5% of the projection-neuron population. In adulthood, however, approximately 22% of projection neurons express Tac1, a substantially higher proportion than that observed at E16.5. One possible explanation is that Tac1 expression is activated later during development in additional projection neurons. Alternatively, developmental changes in Zfhx3 expression could contribute to the differences. However, Zfhx3 expression peaks transiently in dividing cells at E10.5 and declines sharply by E13.5, remaining at comparable levels at P0 and P70 (Osseward et al., 2021), suggesting that it should mark a relatively stable projection-neuron population at both E16.5 and adulthood. The developmental shift we observe is therefore unlikely to be explained by changes in Zfhx3 expression and more likely reflects a genuine change in the population expressing Tac1. Moreover, most dorsal horn neurons are already differentiated and post-migratory at E16.5 as indicated by Ebf1 expression (Hernandez-Miranda et al., 2017). Together, these findings suggest that Tac1 expression is acquired by distinct neuronal populations at different developmental stages, appearing earlier in projection neurons and later in substantial numbers of interneurons.

### Superficial Tac1-lineage neurons predominantly exhibit transient firing properties

Our electrophysiological recordings further revealed substantial heterogeneity among superficial Tac1-lineage neurons. We identified phasic, single-spike, tonic, and reluctant firing patterns, with phasic and single-spike neurons comprising the majority. This differs from the adult Tac1-expressing population characterized by Todd and colleagues, in which delayed firing was prominent (Dickie et al., 2019). In addition to the broader developmental population captured by lineage labeling, this discrepancy may partly reflect differences in laminar distribution, as we primarily targeted neurons in laminae / o, whereas delayed firing is common among excitatory interneurons in lamina (Heinke et al., 2004; Yasaka et al., 2010).

The predominance of phasic firing may have important functional implications. Phasic neurons have been proposed to operate as coincidence detectors, preferentially responding to excitatory inputs arriving within a restricted temporal window rather than continuously integrating inputs over time (Konig et al., 1996; Prescott and De Koninck, 2002, 2005). Such properties could allow superficial Tac1-lineage neurons to respond preferentially to synchronous or rapidly changing sensory inputs. This is particularly intriguing given the majority of primary sensory neurons respond to noxious stimuli (Chisholm et al., 2018; Wang et al., 2018): convergence of inputs on phasic neurons could favor detection of temporally coincident activity associated with noxious events.

Tonic-firing neurons may provide a complementary mode of computation by integrating sustained synaptic input over longer periods (Prescott and De Koninck, 2002, 2005). Their distinct rheobase and action potential properties support functional heterogeneity within the Tac1 lineage, although firing pattern alone cannot identify excitatory, inhibitory, or projection-neuron identity (Lu and Perl, 2005; Todd, 2010; Zheng et al., 2010). Combining electrophysiological characterization with molecular and anatomical identification will therefore be necessary to determine how these intrinsic properties map onto specific Tac1-lineage subpopulations.

### Tac1-lineage neurons are polymodal but encode cooling and heating differently

Our *in vivo* imaging results demonstrate that superficial Tac1-lineage neurons are broadly polymodal but preferentially recruited by noxious stimuli. Approximately 60% responded to mechanical pinch, compared with fewer than 15% responding to innocuous brushing, and neuronal recruitment increased markedly at noxious heating temperatures. Moreover, observations extend previous activity-marker studies showing activation of adult Tac1-expressing neurons by multiple forms of noxious stimulation (Gutierrez-Mecinas et al., 2017) and indicate that Tac1-lineage neurons constitute a convergent sensory population with a strong nociceptive bias.

Interestingly, cooling and heating intensity were represented through distinct coding strategies. Stronger cooling did not substantially recruit additional neurons but instead increased response amplitude within a relative stable cooling-responsive population. In contrast, increasing heat temperature progressively recruited additional neurons, including cells selectively activated at 50°C, without a corresponding increase in response amplitude among broadly heating-responsive neurons. Thus, cooling intensity appears to be represented primarily through amplitude scaling, whereas heating relies more strongly on population recruitment. These findings are consistent with previous *in vivo* imaging of superficial dorsal horn neurons (Ran et al., 2016) and suggest that distinct thermal coding strategies are preserved with the molecularly defined Tac1 lineage.

These imaging results are broadly consistent with a role for Tac1-lineage neurons in nociception. Their strong recruitment by pinch and noxious heat is particularly relevant to the work of Ma and colleagues who showed that ablation of Tac1-Cre;Lbx1-Flpo lineage neurons nearly abolished persistent licking evoked by sustained noxious mechanical, heat, and cold stimulation (Huang et al., 2019). Their findings led to the proposal that spinal Tac1-lineage neurons preferentially contribute to coping response associated with sustained pain. Our functional imaging provides a cellular correlate for this behavioral phenotype: Tac1-lineage neurons are extensively recruited by precisely the types of intense mechanical and thermal stimuli that evoke these coping behaviors.

However, in our experiments, acute optogenetic inhibition of adult spinal Tac1-expressing neurons did not significantly alter licking, shaking, lifting, jumping, or the integrated nociception score during sustained hot- or cold-plate stimulation. This difference from the dramatic phenotype reported after Tac1-lineage ablation raises the possibility that neurons transiently express Tac1 during development contribute importantly to sustained pain-related coping behaviors. Alternatively, the difference may reflect the much stronger and permanent manipulation produced by neuronal ablation compared with acute optogenetic inhibition, or differences in the completeness of neuronal targeting. Thus, rather than contradicting the earlier ablation study, our findings refine its interpretation by suggesting that adult Tac1 expression alone may not define the complete spinal population required for sustained thermal coping behaviors.

## References

Borromeo MD, Meredith DM, Castro DS, Chang JC, Tung KC, Guillemot F, Johnson JE (2014) A transcription factor network specifying inhibitory versus excitatory neurons in the dorsal spinal cord. Development 141:2803–2812.

Braz J, Solorzano C, Wang X, Basbaum AI (2014) Transmitting pain and itch messages: a contemporary view of the spinal cord circuits that generate gate control. Neuron 82:522–536.

Cameron D, Polgar E, Gutierrez-Mecinas M, Gomez-Lima M, Watanabe M, Todd AJ (2015) The organisation of spinoparabrachial neurons in the mouse. Pain 156:2061–2071.

Chisholm KI, Khovanov N, Lopes DM, La Russa F, McMahon SB (2018) Large Scale In Vivo Recording of Sensory Neuron Activity with GCaMP6. eNeuro 5.

Choi S, Hachisuka J, Brett MA, Magee AR, Omori Y, Iqbal NU, Zhang D, DeLisle MM, Wolfson RL, Bai L, Santiago C, Gong S, Goulding M, Heintz N, Koerber HR, Ross SE, Ginty DD (2020) Parallel ascending spinal pathways for affective touch and pain. Nature 587:258–263.

Das Gupta RR, Scheurer L, Pelczar P, Wildner H, Zeilhofer HU (2021) Neuron-specific spinal cord translatomes reveal a neuropeptide code for mouse dorsal horn excitatory neurons. Sci Rep 11:5232.

Davis OC, Dickie AC, Mustapa MB, Boyle KA, Browne TJ, Gradwell MA, Smith KM, Polgar E, Bell AM, Kokai E, Watanabe M, Wildner H, Zeilhofer HU, Ginty DD, Callister RJ, Graham BA, Todd AJ, Hughes DI (2023) Calretinin-expressing islet cells are a source of pre- and post-synaptic inhibition of non-peptidergic nociceptor input to the mouse spinal cord. Sci Rep 13:11561.

Dickie AC, Bell AM, Iwagaki N, Polgar E, Gutierrez-Mecinas M, Kelly R, Lyon H, Turnbull K, West SJ, Etlin A, Braz J, Watanabe M, Bennett DLH, Basbaum AI, Riddell JS, Todd AJ (2019) Morphological and functional properties distinguish the substance P and gastrin-releasing peptide subsets of excitatory interneuron in the spinal cord dorsal horn. Pain 160:442–462.

Ding YQ, Yin J, Kania A, Zhao ZQ, Johnson RL, Chen ZF (2004) Lmx1b controls the differentiation and migration of the superficial dorsal horn neurons of the spinal cord. Development 131:3693–3703.

Garel S, Marin F, Grosschedl R, Charnay P (1999) Ebf1 controls early cell differentiation in the embryonic striatum. Development 126:5285–5294.

Graham BA, Hughes DI (2020) Defining populations of dorsal horn interneurons. Pain 161:2434–2436.

Grudt TJ, Perl ER (2002) Correlations between neuronal morphology and electrophysiological features in the rodent superficial dorsal horn. J Physiol 540:189–207.

Gutierrez-Mecinas M, Polgar E, Bell AM, Herau M, Todd AJ (2018) Substance P-expressing excitatory interneurons in the mouse superficial dorsal horn provide a propriospinal input to the lateral spinal nucleus. Brain Struct Funct 223:2377–2392.

Gutierrez-Mecinas M, Bell AM, Marin A, Taylor R, Boyle KA, Furuta T, Watanabe M, Polgar E, Todd AJ (2017) Preprotachykinin A is expressed by a distinct population of excitatory neurons in the mouse superficial spinal dorsal horn including cells that respond to noxious and pruritic stimuli. Pain 158:440–456.

Han X, Chow BY, Zhou H, Klapoetke NC, Chuong A, Rajimehr R, Yang A, Baratta MV, Winkle J, Desimone R, Boyden ES (2011) A high-light sensitivity optical neural silencer: development and application to optogenetic control of non-human primate cortex. Front Syst Neurosci 5:18.

Heinke B, Ruscheweyh R, Forsthuber L, Wunderbaldinger G, Sandkuhler J (2004) Physiological, neurochemical and morphological properties of a subgroup of GABAergic spinal lamina II neurones identified by expression of green fluorescent protein in mice. J Physiol 560:249–266.

Hernandez-Miranda LR, Muller T, Birchmeier C (2017) The dorsal spinal cord and hindbrain: From developmental mechanisms to functional circuits. Dev Biol 432:34–42.

Huang T, Lin SH, Malewicz NM, Zhang Y, Zhang Y, Goulding M, LaMotte RH, Ma Q (2019) Identifying the pathways required for coping behaviours associated with sustained pain. Nature 565:86–90.

Konig P, Engel AK, Singer W (1996) Integrator or coincidence detector? The role of the cortical neuron revisited. Trends Neurosci 19:130–137.

Larsson M (2017) Pax2 is persistently expressed by GABAergic neurons throughout the adult rat dorsal horn. Neurosci Lett 638:96–101.

Littlewood NK, Todd AJ, Spike RC, Watt C, Shehab SA (1995) The types of neuron in spinal dorsal horn which possess neurokinin-1 receptors. Neuroscience 66:597–608.

Lu Y, Perl ER (2005) Modular organization of excitatory circuits between neurons of the spinal superficial dorsal horn (laminae I and II). J Neurosci 25:3900–3907.

Ma Q (2022) A functional subdivision within the somatosensory system and its implications for pain research. Neuron 110:749–769.

Mo A, Mukamel EA, Davis FP, Luo C, Henry GL, Picard S, Urich MA, Nery JR, Sejnowski TJ, Lister R, Eddy SR, Ecker JR, Nathans J (2015) Epigenomic Signatures of Neuronal Diversity in the Mammalian Brain. Neuron 86:1369–1384.

Osseward PJ, 2nd, Amin ND, Moore JD, Temple BA, Barriga BK, Bachmann LC, Beltran F, Jr., Gullo M, Clark RC, Driscoll SP, Pfaff SL, Hayashi M (2021) Conserved genetic signatures parcellate cardinal spinal neuron classes into local and projection subsets. Science 372:385–393.

Polgar E, Bell AM, Gutierrez-Mecinas M, Dickie AC, Akar O, Costreie M, Watanabe M, Todd AJ (2020) Substance P-expressing Neurons in the Superficial Dorsal Horn of the Mouse Spinal Cord: Insights into Their Functions and their Roles in Synaptic Circuits. Neuroscience 450:113–125.

Prescott SA, De Koninck Y (2002) Four cell types with distinctive membrane properties and morphologies in lamina I of the spinal dorsal horn of the adult rat. J Physiol 539:817–836.

Prescott SA, De Koninck Y (2005) Integration time in a subset of spinal lamina I neurons is lengthened by sodium and calcium currents acting synergistically to prolong subthreshold depolarization. J Neurosci 25:4743–4754.

Ran C, Hoon MA, Chen X (2016) The coding of cutaneous temperature in the spinal cord. Nat Neurosci 19:1201–1209.

Ribeiro-da-Silva A, Hokfelt T (2000) Neuroanatomical localisation of Substance P in the CNS and sensory neurons. Neuropeptides 34:256–271.

Roome RB, Bourojeni FB, Mona B, Rastegar-Pouyani S, Blain R, Dumouchel A, Salesse C, Thompson WS, Brookbank M, Gitton Y, Tessarollo L, Goulding M, Johnson JE, Kmita M, Chedotal A, Kania A (2020) Phox2a Defines a Developmental Origin of the Anterolateral System in Mice and Humans. Cell Rep 33:108425.

Todd AJ (2010) Neuronal circuitry for pain processing in the dorsal horn. Nat Rev Neurosci 11:823–836.

Vrontou S, Wong AM, Rau KK, Koerber HR, Anderson DJ (2013) Genetic identification of C fibres that detect massage-like stroking of hairy skin in vivo. Nature 493:669–673.

Wang F, Belanger E, Cote SL, Desrosiers P, Prescott SA, Cote DC, De Koninck Y (2018) Sensory Afferents Use Different Coding Strategies for Heat and Cold. Cell Rep 23:2001–2013.

Wang LH, Ding WQ, Sun YG (2022) Spinal ascending pathways for somatosensory information processing. Trends Neurosci 45:594–607.

Wercberger R, Basbaum AI (2019) Spinal cord projection neurons: a superficial, and also deep, analysis. Curr Opin Physiol 11:109–115.

Xu Y, Lopes C, Wende H, Guo Z, Cheng L, Birchmeier C, Ma Q (2013) Ontogeny of excitatory spinal neurons processing distinct somatic sensory modalities. J Neurosci 33:14738–14748.

Yasaka T, Tiong SYX, Hughes DI, Riddell JS, Todd AJ (2010) Populations of inhibitory and excitatory interneurons in lamina II of the adult rat spinal dorsal horn revealed by a combined electrophysiological and anatomical approach. Pain 151:475–488.

Zheng J, Lu Y, Perl ER (2010) Inhibitory neurones of the spinal substantia gelatinosa mediate interaction of signals from primary afferents. J Physiol 588:2065–2075.

