## Supplemental Material for "Superficial spinal Tac1-lineage neurons are polymodal nociceptive"

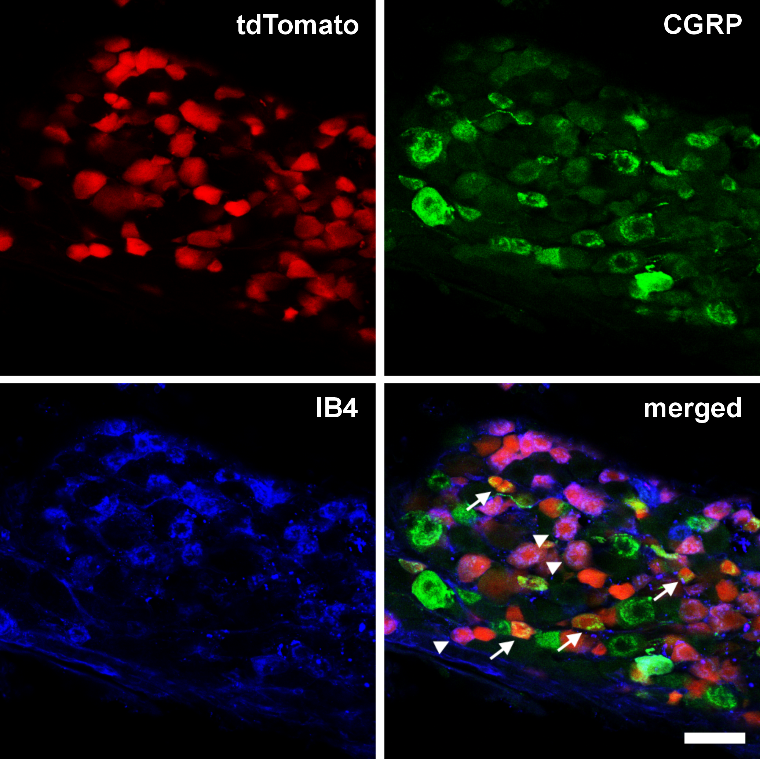

**Figure S1. Tac1 is expressed in dorsal root ganglion neurons**

Representative DRG section from a *Tac1-tdTomato* mouse showing tdTomato (red), CGRP (green), and IB4 (blue) labeling. Scale bars = 50 μm.

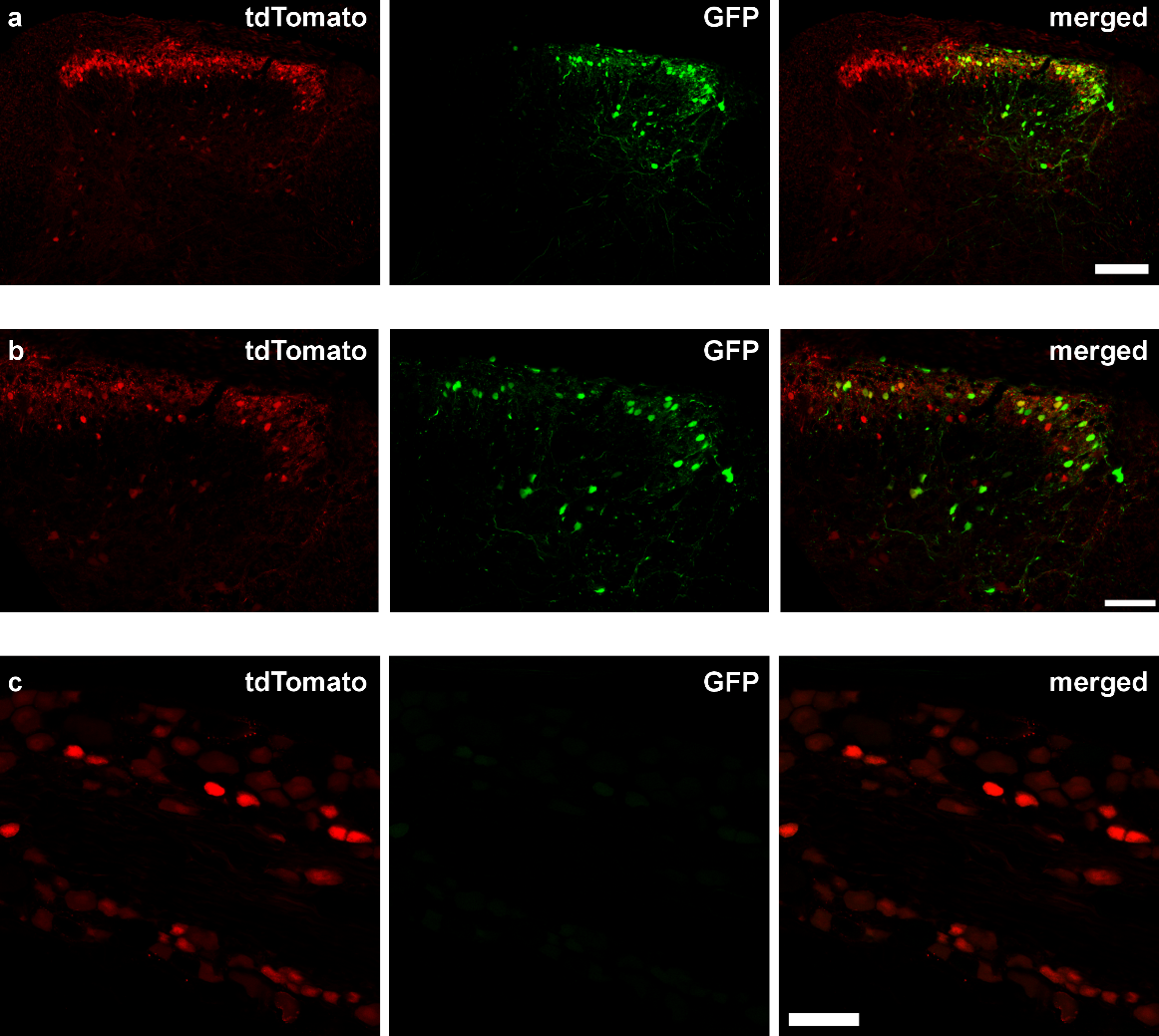

**Figure S2. Intraspinal viral injection did not transfect DRG neurons**

(a) Representative spinal cord section from a *Tac1-tdTomato* mouse following intraspinal injection of an AAV carrying Cre-dependent GFP, showing GFP (green) and tdTomato (red) expression in the dorsal horn.
(b) Higher-magnification images of the same spinal cord section shown in (a).
(c) The injected AAV did not transfect DRG neurons. Scale bars = 100 μm (a) and 50 μm (b and c).

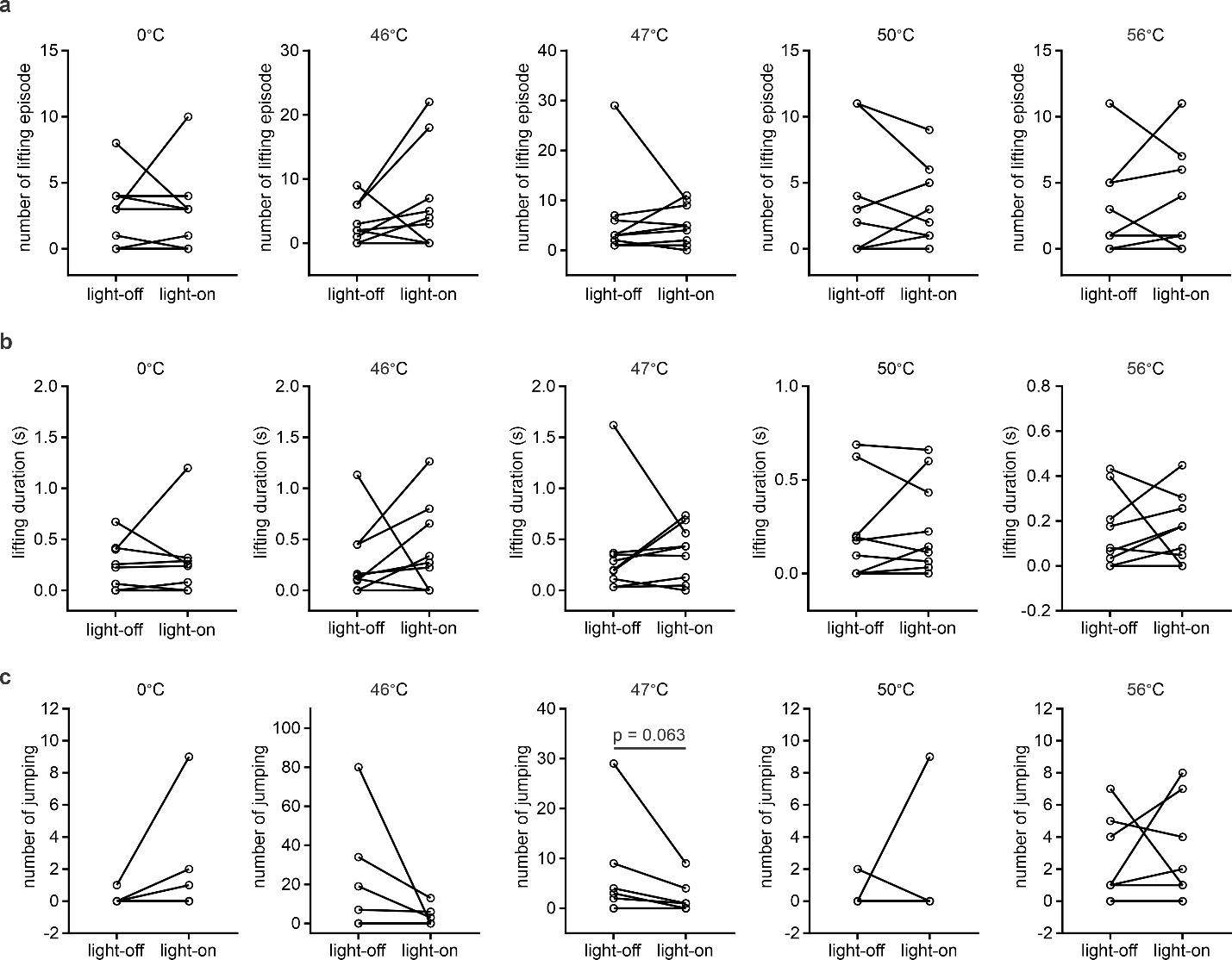

**Figure S3. Adult spinal Tac1-expressing neurons are not required for coping behaviors evoked by sustained noxious thermal stimuli**

Optogenetic inhibition of spinal Tac1-expressing neurons did not significantly alter the number of lifting episodes (a), lifting duration (b), or number of jumping (c). Statistical comparisons were performed using Wilcoxon matched-pairs signed-rank test. n = 9 *Tac1-Cre* mice.

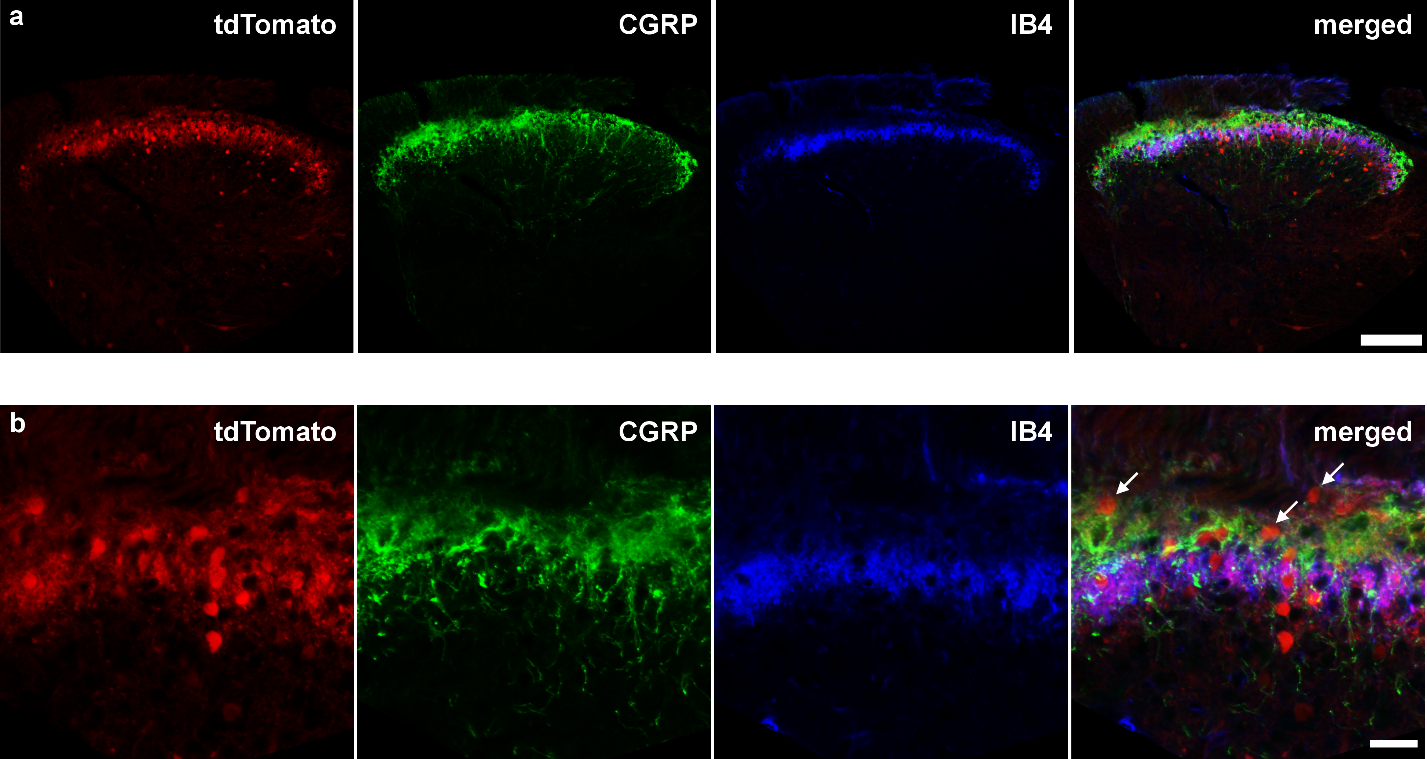

**Figure S4. Distribution of Tac1-lineage neurons in adult dorsal horn**

(a) Representative spinal cord section from a *Tac1-tdTomato* mouse showing tdTomato (red), CGRP (green), and IB4 (blue) labeling in the dorsal horn.
(b) Higher-magnification images of the same spinal cord section shown in (a). Tac1-lineage neurons are distributed across multiple laminae of the dorsal horn, with extensive tdTomato labeling in primary sensory afferents. Scale bars = 100 μm (a) and 50 μm (b).

**Table 1. Antibodies and dyes used for immunostaining**

| Antibodies or dye | Sources | Dilution |
| --- | --- | --- |
| Goat anti-hPax2 | AF3364, R&D Systems | 1:1,000 |
| Guinea pig anti-Lmx1b | Gift from Carmen Birchmeier (Max Delbruck Center, Berlin) | 1:10,000 |
| Rabbit anti-Ebf1 | PA5-13540, ThermoFisher | 1:100 |
| Mouse anti-GFP | ab1218, Abcam | 1:500 |
| Sheep anti Zfhx3 | AF7384, R&D Systems | 1:50 |
| Rabbit anti-CGRP | C8198, Sigma | 1:1000 |
| IB4 Alexa-647 | I32450, Invitrogen | 1:500 |
